# COVID-19 Biomarkers in research: Extension of the OncoMX cancer biomarker data model to capture biomarker data from other diseases

**DOI:** 10.1101/2020.09.09.196220

**Authors:** N Gogate, D Lyman, K.A Crandall, R Kahsay, D.A Natale, S Sen, R Mazumder

## Abstract

Scientists, medical researchers, and health care workers have mobilized worldwide in response to the outbreak of COVID-19, caused by severe acute respiratory syndrome coronavirus 2 (SARS-CoV-2; SCoV2). Preliminary data have captured a wide range of host responses, symptoms, and lingering problems post-recovery within the human population. These variable clinical manifestations suggest differences in influential factors, such as innate and adaptive host immunity, existing or underlying health conditions, co-morbidities, genetics, and other factors. As COVID-19-related data continue to accumulate from disparate groups, the heterogeneous nature of these datasets poses challenges for efficient extrapolation of meaningful observations, hindering translation of information into clinical applications. Attempts to utilize, analyze, or combine biomarker datasets from multiple sources have shown to be inefficient and complicated, without a unifying resource. As such, there is an urgent need within the research community for the rapid development of an integrated and harmonized COVID-19 Biomarker Knowledgebase. By leveraging data collection and integration methods, backed by a robust data model developed to capture cancer biomarker data we have rapidly crowdsourced the collection and harmonization of COVID-19 biomarkers. Our resource currently has 138 unique biomarkers. We found multiple instances of the same biomarker substance being suggested as multiple biomarker types during our extensive cross-validation and manual curation. As a result, our Knowledgebase currently has 265 biomarker type combinations. Every biomarker entry is made comprehensive by bringing in together ancillary data from multiple sources such as biomarker accessions (canonical UniProtKB accession, PubChem Compound ID, Cell Ontology ID, Protein Ontology ID, NCI Thesaurus Code, and Disease Ontology ID), BEST biomarker category, and specimen type (Uberon Anatomy Ontology) unified with ontology standards. Our preliminary observations show distinct trends in the collated biomarkers. Most biomarkers are related to the immune system (SAA,TNF-∝, and IP-10) or coagulopathies (D-dimer, antithrombin, and VWF) and a few have already been established as cancer biomarkers (ACE2, IL-6, IL-4 and IL-2). These trends align with proposed hypotheses of clinical manifestations compounding the complexity of COVID-19 pathobiology. We explore these trends as we put forth a COVID-19 biomarker resource that will help researchers and diagnosticians alike. All biomarker data are freely available from https://data.oncomx.org/covid19.

## Introduction

The devastating outbreak of the novel, highly contagious Coronavirus Disease (COVID-19), originating in Wuhan, China, has rapidly spread worldwide since first reported to the world in early January 2020. The genomic sequence of the causative betacoronavirus, SARS-CoV-2 (Severe Acute Respiratory Syndrome Coronavirus 2; SCoV2), was first released on January 7, 2020 with similarity to SCoV (Severe Acute Respiratory Syndrome Coronavirus) and MERS-CoV (Middle East Respiratory Syndrome Coronavirus)^1–5^. COVID-19 has created major challenges for worldwide health systems, caused global disruption, and far-reaching consequences to the global economy^6, 7^. In response, the World Health Organization (WHO) declared a global pandemic; as of August 31st 2020, there are more than 25 million confirmed cases globally and more than 840,000 reported fatalities5, ^8–10^. While researchers race to find a drug(s) or vaccine(s) for the virus, a critical need to identify biomarkers for COVID-19 disease has become evident. A simple search in Google Scholar for `COVID-19 biomarkers` retrieves more than 10,000 records since 2020. Not all of these publications describe a biomarker, but many refer to them. Preliminary evaluation reveals that almost none of the biomarker data described in these references are standardized and harmonized to existing ontologies and terms.

SCoV2 is the seventh coronavirus known to infect humans^11^. Coronaviruses SCoV, MERS-CoV, and SCoV2 can cause severe disease; while Coronaviruses HKU1, NL63, OC43, and 229E are associated with mild disease states^12, 13^. High recombination rates and genetic diversity of coronaviruses in the wild suggest that further outbreaks and unpredictable virulence will likely arise in future recombinants^14^, which could lead to different outcomes in patients. Serious clinical manifestations of COVID-19 (in some individuals) include: severe acute respiratory syndrome, inflammatory pneumonitis, hypoxia, blood clots, embolisms, gastrointestinal illness, cardiac and vascular damage, and organ damage (lung, heart, kidney, liver, brain)^3^. Severity and mortality of COVID-19 appear to be more prevalent in men (3.1/million) than women (2.7/million) and, overall more so in the elderly with underlying health conditions, such as hypertension, cardiovascular disease, immunosenescence, immunocompromised systems, and diabetes^3, 5, 14–16^. Clinical observations of hospitalized COVID-19 patients report lymphopenia and monocytopenia, , and hypoalbuminemia, as well as elevated proinflammatory cytokines (“cytokine storm”). In severe cases, pneumonia with a “ground glass” opacity in chest CT scans, lung injury, and pneumonitis are typically observed^8, 17, 18^. Lymphopenia and the cytokine storm may initiate severe COVID-19 pathogenesis, viral sepsis, inflammatory-induced lung injury and pneumonitis, acute respiratory distress syndrome (ARDS), respiratory failure, shock, organ failure, and death^3, 19, 20^. Probability of the severe damage from the direct or indirect effects of SARS-CoV-2 replication can be exacerbated by underlaying injury caused by chronic conditions like hypertension, diabetes and/or cancer^16, 21, 22^. Identification of physiological or pathological differences associated with poor outcomes of the COVID-19 in patients with underlying conditions and discovery of prospective biomarkers predictive of these outcomes is of paramount importance.

The global impact of COVID-19 has mobilized the biomedical community – from bench to bed – to combat the pathogen. Scientific and clinical observations have accrued in dispersed resources in the effort to publicize the data as rapidly as possible for investigation. Representative features of COVID-19 have therefore begun to surface, but deeper <u>reproducible measures (biomarkers) of COVID-19 pathobiology and pharmacologic intervention have yet to emerge</u>. Experience suggests that significant data for **nucleic acid, protein, glycan, and other biomarker material await discovery by cross-disciplinary investigations of COVID-19 publications and repositories**. While preliminary discoveries demonstrate clinical applicability of potential biomarkers, additional research must establish specificity and sensitivity during risk assessment, diagnostic measurements, or therapeutic applications to a particular disease state^23, 24^. For both research and clinical applications, improved methods for aggregation of biomarker knowledge must be implemented, a process that comparatively lags behind due to the heterogeneous nature of biomarker data. As a result, **improving the methods by which we explore, identify, and discover biomarkers is a vital and necessary focus of biomedical research**. We provide for immediate use a publicly available compilation of COVID-19 biomarkers that enables academic and regulatory scientists along with industry researchers to explore up to date COVID-19 biomarkers in different stages of development and application.

### Biomarker Research and Data Integration Challenges

The FDA-NIH Biomarker Working Group (FNBWG) defines a biomarker as a “characteristic that is measured as an indicator of normal biological processes, pathogenic processes, or responses to an exposure or intervention, including therapeutic interventions”^25^. Molecular biomarkers (also known as molecular markers or signature molecules) may be genes, proteins, glycans, or metabolites, for example, that may be used in different stages of disease assessment and treatment evaluation, but differ from clinical assessments and have distinct functions in biomedical research, clinical practice, and medical product development. The FNBWG further distinguishes important subtypes, by role, for which some instances have been identified as potential COVID-19 markers (Fig. 1): **Diagnostic^26, 27^**, **Monitoring^28^**, **Pharmacodynamic/Response ^29^**, **Predictive^30^**, **Prognostic^31^**, **Safety^32^**, and **Susceptibility/Risk^26,33^**. Measured as objective, reproducible numeric or categoric values, biomarkers play a significant role in highlighting the relationships among environmental exposures, human biology, and disease^34^. The BEST Resource provides a constructive framework by which to organize, standardize, and integrate data elements in the COVID-19 biomarker resource.

**Figure 1.**
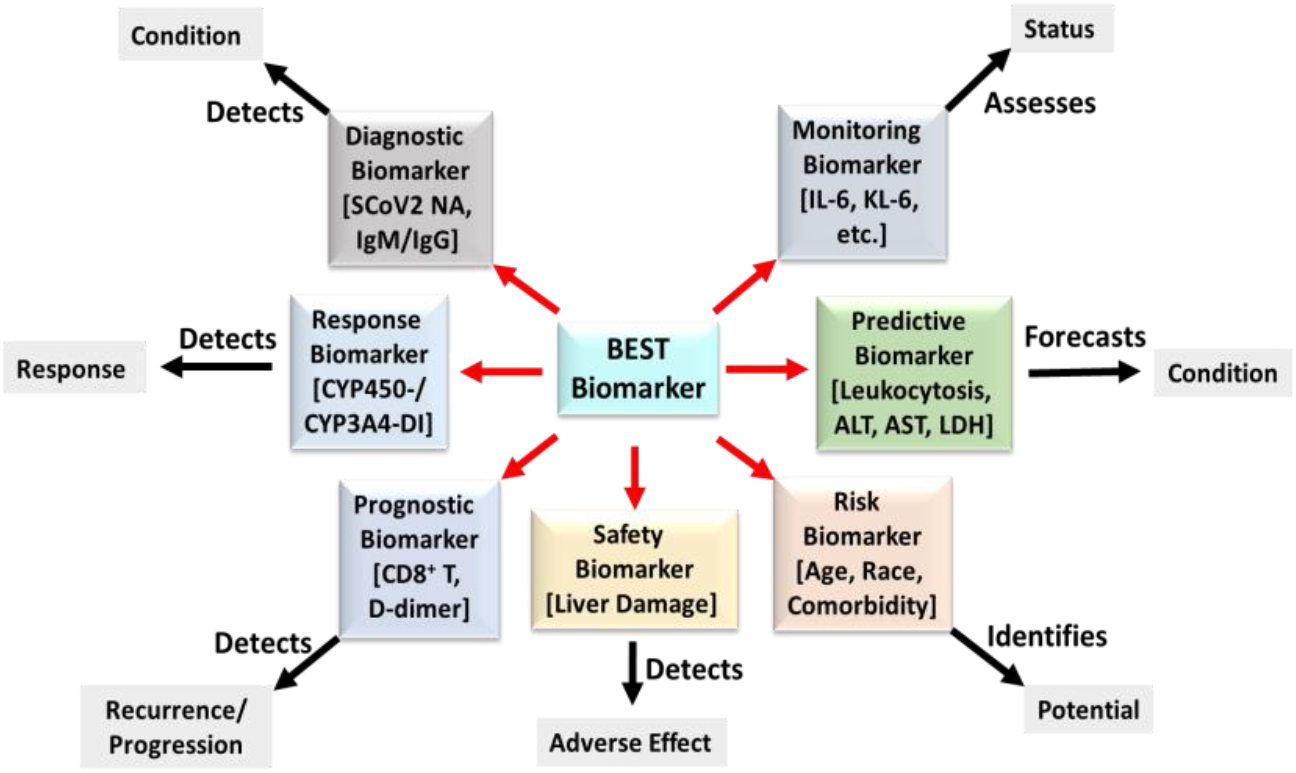
Model of BEST biomarker subtypes. Each category of BEST Biomarker, indicated by red arrows, fulfills a distinct role “as an indicator of normal biological processes, pathogenic processes, or responses to an exposure or intervention.” Tests for specific instances of potential COVID-19 biomarkers (in brackets) provide measurable evidence data of existing or potential health status.

### Standards and Ontologies

The biomarker list is created for easy integration in other biomarker knowledgebases that use standards and ontologies such as our recently developed infrastructure (OncoMX)^35^, a biomarker knowledgebase designed to integrate cancer-centric data and combine it with newly generated data, establishing a strong application customized for biomarker research. This platform implements strict adherence to accepted standards used in major resources, such as National Center for Biotechnology Information (NCBI)^36^, European Bioinformatics Institute (EBI)^37^, Alliance of Genome Resources^38^, and others. These resources, and OncoMX, rely heavily on existing and new biomedical standards and ontologies for semantic unification of datasets, which can enable efficient knowledge modeling, information retrieval, and data sharing across otherwise diverse data^39–43^. The emphasis on leveraging existing standards and ontologies promotes extensibility and sustainability, allowing the platform to focus on data quality, integration, standardization, and knowledgebase maintenance and extension. The Uberon Anatomy Ontology^44^ and other ontologies are employed in OncoMX, for example, to support efficient cross comparison of biomarker data of human and model organism orthologs.

## Materials and Methods

The goal of the project is to collect and harmonize COVID-19 biomarkers that will assist researchers working on the development of diagnostics or drugs by organizing biomarker data from publications and bioinformatic databases into a standardized table. Various steps - from data collection, organization, standardization, and integration of the data elements - are involved to bring together the COVID-19 biomarkers (as depicted in Fig. 2).

**Figure 2.**
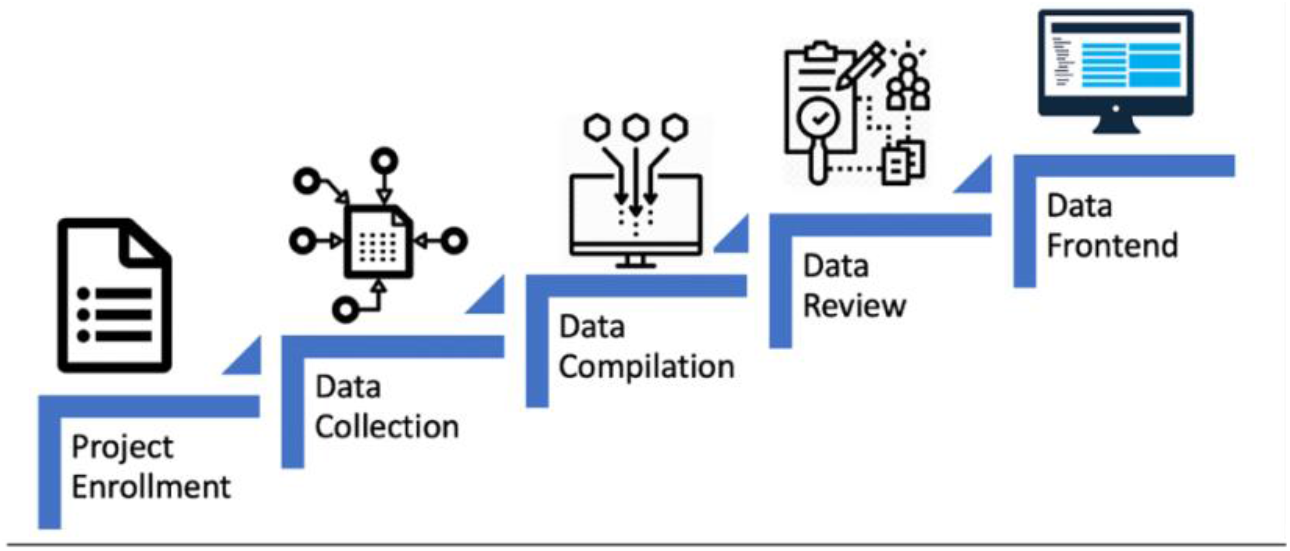
Steps for collection, organization, standardization and integration of the data elements in the COVID-19 biomarker resource data model.

### Crowdsourcing

With the exponential rise in the number of publications and publication drafts on COVID-19 biomarkers it was clear that a crowdsourcing effort using a robust data model was needed to rapidly collect and cross-validate the data. The project was advertised to collaborating faculty members which led to the recruitment of approximately 30 volunteers. Key staff members of the OncoMX team led the volunteers and served as reviewers for all annotations. The groups were organized into reviewers and curators. Curators read publications and filled in tables. Reviewers then read the publications and ensured that the cell entries were correct prior to moving them to the reviewed biomarker table.

### Data Collection and Compilation

Curators searched for “COVID-19 biomarkers” in Google Scholar that were publicly available after January 2020. Information about a biomarker and its role in COVID-19 was retrieved from selected articles and filled into a structured format. The data collected were mainly populated into columns like biomarker name, measured biomarker, specimen type, biomarker description and drug mentioned. Ancillary data regarding the biomarker such as biomarker accession and BEST biomarker type were further mapped to each biomarker entry. Biomarker accession could include the canonical UniProtKB accession^45^, PubChem Compound ID^46^, Cell Ontology ID^47^, Protein Ontology ID^48^, and Disease Ontology ID^49^. The curator could write notes in a free text column that documented any comments regarding the data curated. Each curator would upload their file into a shared drive once every week. Reviewers would then compile the data from all the curators into a single cohesive dataset file marked Unreviewed.

### Review Processing and Quality Check

The unreviewed data file was then scrutinized by at least two reviewers with experience in curation of biomarkers and ontology mapping. All annotations were checked for content. These checks included: confirmation of biomarker name mentioned in the article, appropriate mapping of the biomarker accession, suitable representation of the BEST biomarker type based on the article and definitions provided by the FNBWG, and documentation of the specimen type with mapping to Uberon anatomical IDs. Table 1 provides details on the rubric of approved data types in each of the columns. Curator notes were carefully assessed during reviews of the biomarker entries to answer any queries that arose while curating the data. The data points were checked for completeness and adherence to the rubric of data collection (Table 1). Resulting entries were compiled and quality checked to ensure integrity and format stability.

**Table 1.** Biomarker table header descriptions and column content.

| Table 1. Biomarker table header descriptions and column content. |  |
| --- | --- |
| Header | Column Content |
| literature_evidence | PMID (PubMed ID) or DOI (DOI number) |
| biomarker_substance_accession | UPKB (UniProtKB/Swiss-Prot ac) or PCCID (PubChem Compound ID) or CO (Cell Ontology ID) or PRO (Protein Ontology ID) or DO (Disease Ontology ID) |
| biomarker_name | Common name or UniProtKB protein name and gene symbol or short name in parenthesis |
| measured_biomarker | increased/decreased level or expression or counts or ratio |
| specimen_type | Uberon name (Uberon ID) |
| BEST_biomarker_type | monitoring; diagnostic; prognostic; predictive; risk/susceptibility; safety; pharmacodynamic/response |
| drug | Drug name (DrugBank ID) |
| biomarker_description | Free text from paper (preferably unedited) |
| curator_name | First and Last name |
| curator_ORCID | ORCID |
| curator_notes | Free text from curator |
| reviewer_initial | Name/Initial: Free text from reviewer |

## Results and Discussion

The first phase of the project lasted approximately six weeks (June 22 – July 31, 2020). Based on usefulness of the collected biomarkers and availability of volunteers the cycle will be repeated monthly. The data model for the COVID-19 biomarkers is based on the OncoMX cancer biomarker model.

Crowdsourcing allowed us to annotate and cross-validate 265 biomarker type combinations. In this effort, we have found a number of issues. For example, we note that the same biomarker is often known by multiple different names in the domain of biomarkers, which might differ from the name used in standardized databases - thus potentially making the connection between biomarker and underlying biology less amenable to discovery. One such case is Carbohydrate Antigen 15-3, also commonly known as Krebs von den Lungen-6, which is called Mucin-1 in UniProtKB. Our knowledgebase has solved these discrepancies by including all such entries identified by respective studies, and then unifying them under common identifier links to standardized databases. Another issue involves discerning exactly what is being measured. For example, what substance is being assayed when using alanine aminotransferase (ALT) as a biomarker? Is it alanine aminotransferase 1, alanine aminotransferase 2, or both? It is both. Finally, there is a chance that a biomarker obtained from one source could be diagnostic for a particular disease, but that same biomarker obtained from a different source might not be. For example, soluble urokinase plasminogen activator receptor (suPAR) isolated from ovarian cysts appears to distinguish between malignant vs benign cysts^50^, but that same substance is only prognostic (for a number of diseases) when obtained from blood^51^. Such common issues, often faced while compiling of a knowledgebase based on an expansive literature, have been addressed in our resource.

A total of 138 biomarkers were corroborated, and classified into the diagnostic, monitoring, prognostic, predictive, risk, response and safety categories for COVID-19.The top few biomarkers in terms of the number of manuscripts in which they were mentioned were C-reactive protein (CRP) (41), Interleukin-6 (IL-6) (24), D-dimer (22), neutrophil to lymphocyte ratio (NLR) (15), serum amyloid A (SAA) (13) lymphocyte count (10), CD4+ counts (8), and CD8+ counts (8) (Fig. 3). Additionally, we also identified multiple biomarkers rarely mentioned in connection with COVID-19. Antithrombin, von Willebrand factor (VWF), Citrullinated H3 (Cit-H3), macrophage colony stimulating factor (MCSF), and Ewing Sarcoma RNA binding protein (EWS) were leading rare biomarkers that showed potential for further investigation. Our results indicate the emergence of a pattern that points to specific pathways and cell types targeted by SARS-CoV-2. Our manually curated resource shows that most biomarkers belong to these biological processes within specific tissue systems and supports the idea that further investigations using multidrug combination therapies targeting these biological processes, is needed.

**Figure 3.**
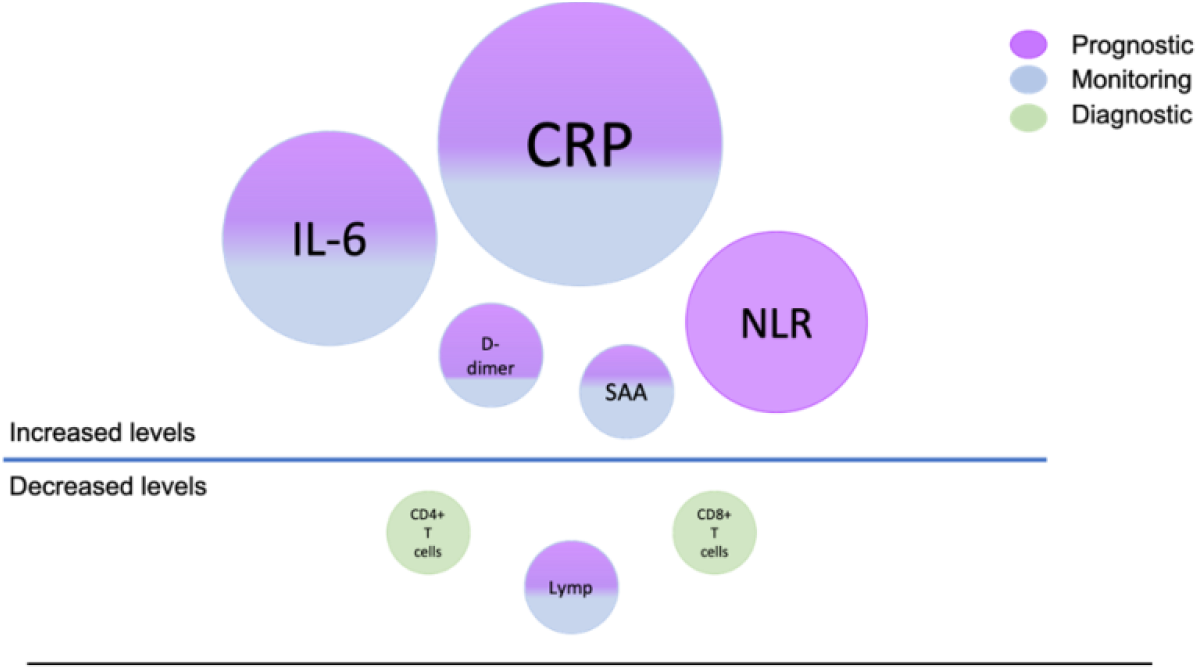
COVID-19 Biomarkers highlights. Top biomarkers are depicted as increased or decreased levels. The size of the circle is indicative of the number of articles supporting the biomarker. The color of the circle corresponds to the BEST biomarker type (Blue – monitoring, Purple – prognostic, Green – diagnostic).

CRP was identified as a top biomarker, appearing in forty-one independent studies. A high level of CRP indicates disease progression and has been positively correlated with lung lesions^52^. It has been used as a monitoring biomarker in early stages of the disease to determine progression from mild to severe ^52, 53^. IL-6 expression is high in lung and arteries for healthy individuals^54^. IL-6 is a known cancer biomarker^35^. It was also used as a disease progression monitoring biomarker in multiple studies and its elevated levels were shown to be strongly associated with respiratory failure in symptomatic COVID-19 patients. Diagnosticians have used this biomarker to determine the need for mechanical ventilation^55^. Other interleukins, such as IL-4 and IL-2, were also proposed in some studies. Of note, interleukins can be both pro and anti-inflammatory, and as such, use of these biomarkers should be coupled with other indicators of disease progression. Interestingly, Herold *et al.* found no correlation between IL-6 levels and age, comorbidities, radiological findings, respiratory rate, or qSofa score of patients^55^; on the other hand, IL-4 was shown to inhibit SARS-CoV replication partially through down regulation of ACE2 expression *in vitro*^56^. While this study investigated SARS-CoV and not SARS-CoV-2, research advises screening of all patients for hyper inflammation^57^ and the use of inflammation biomarkers to assess severity of disease. NLR was also suggested as a prognostic biomarker in over 10 independent studies. It has been commonly used as a marker for subclinical or systemic inflammation and a high NLR has been linked to poor clinical outcome in many solid tumors. For COVID-19 patients of advanced age, this ratio, if elevated, should serve as a prognostic biomarker to determine access to valuable limited clinical resources like intensive care units (ICUs) or ventilators^58^. Another biomarker reported by multiple research groups, in agreement with aggravated inflammation observed in COVID-19 patients, is SAA. SAA was used as both a monitoring and a prognostic biomarker by multiple independent research groups to evaluate severity and prognosis of COVID-19. Dynamic changes in SAA have been proposed as prognostic markers in COVID-19 progression^59^. This is because it belongs to the family of apolipoproteins that are constitutively expressed in plasma. SAA is a potential therapeutic target in chronic inflammation^60^. MCSF, another biomarker that plays an important role in macrophage homeostasis, and has been proposed as a therapeutic target that warrants further analysis^61, 62^. A common theme from our resource points to the immune system of the patients and the inflammatory response to the disease. Other concurrent biomarkers in our resource are tumor necrosis factor-∝ (TNF-∝), interferon-ϒ inducible protein-10 (IP-10), CD4+ counts and CD8+ counts, all of which are suggestive of the immune system of the patient as a central target to determine disease progression, therapy, and possibly, prevention.

Another biomarker that has been extensively studied is D-dimer which is a degradation product of crosslinked fibrin resulting from plasmin cleavage^63^. Several independent studies from Wuhan, China, have shown that elevated levels of D-dimer in COVID-19 patients are associated with higher mortality. Indeed, it has been used as a prognostic biomarker to predict mortality rates in patients with COVID-19^64^. However, since it is a product of cross-linked fibrin there are many other common conditions in which it can be elevated and, consequently, use of this biomarker warrants caution. The most common substances resulting in analytical interference with D-dimer levels are paraproteins, bilirubin, lipids, and hemolysis^63^. As such, establishing a fold-change cutoff specific for the patient for this prognostic biomarker before drawing conclusions is essential. Interestingly, our resource suggests the use of other biomarkers from similar biological processes, though these have been studied less extensively. These include VWF and antithrombin, among others. VWF, made within endothelial cells, helps platelets stick together, assists clot formation, and transports coagulation factor VIII to areas of clot formation^65^. High levels of VWF have been linked to potential efficacy of COVID-19 treatment and have also been used as a prognostic biomarker for endothelial damage^66^. Escher *et al.* observed an approximate 500% increase in VWF and coagulation factor VIII expression in COVID-19 patients during later stages of stay in an intensive care unit (ICU). This increase was observed in patients that registered an increase in D-dimer expression in the earlier stages of their stay in the ICU. These patients underwent extensive endothelial stimulation and damage, which can be explained by the presence of ACE2, the receptor for SARS-CoV-2, on the surface of endothelial cells^67^. Similarly, antithrombin, a glycoprotein that plays a critical role in controlling coagulation^68^, was also proposed as a biomarker by at least 2 independent studies. Decreased levels of antithrombin, along with increased levels of D-dimer and VWF, in conjunction with other proposed biomarkers like fibrinogen expression and platelet counts, point to recurrent coagulopathies in COVID-19 patients.

A biomarker that seems to have given interesting results, in the context of metabolic syndrome, diabetes, and hyperlipidemia, is Low Density Lipoprotein (LDL)^69–71^. Subjects with pre-existing hyperlipidemia appear to have decreased in LDL at the onset of the disease, with lower levels predicting worse outcome^69^. It is possible that virus particles require LDL to proliferate and replicate. The drop in LDL at the onset indicates huge virus reproduction capability, rather than a good indicator in the context of metabolic syndrome. However, even in these subjects, lowering of endogenous LDL production with medications such as statin may help to reduce or at least impair virus reproduction capability^69^.

Finally, our resource has also registered biomarkers like EWS, Cit-H3, and ACE2, that have extensive ramifications in various cancers. EWS, primarily implicated in Ewing’s Sarcoma, plays a role in transcriptional repression, and promotes tumorigenesis by forming fusion proteins^72, 73^. Cit-H3 plays an important role in neutrophil release of nuclear chromatin, also called neutrophil extracellular traps (NETs), which have been associated with tumor progression in colon cancer^74, 75^. Notably, NETs have also been proposed as a biomarker for SARS-CoV-2 infection in COVID-19 patients and is known to play a role in thrombosis, thereby strengthening our observation that specific biological processes are activated during SARS-CoV-2 infection. Finally, the angiotensin converting enzyme ACE2 that serves as a receptor for the spike glycoprotein of SARS-CoV-2, shows stabilized protein levels in colorectal and renal cancers, and has also been proposed to be used as a biomarker^76^. This suggests that the majority of cancer patients, and not just immunocompromised patients, also have an elevated risk of contracting the disease, and might have a poorer prognosis when compared to non-cancer individuals with COVID-19.

Taken together, these data suggest that hyper-activation of the immune system, coagulopathies and the targeting of specific types of cells that are indispensable for vasculature like the endothelial cells are the primary *modus operandi* of SARS-CoV-2, and combination therapies targeting these biological processes may be beneficial for COVID-19 patients. Additionally, risk biomarkers like cardiovascular disease, hypertension, thrombocytopenia, and cancer, need to be taken into consideration while devising therapeutic regimens. Our manually curated and regularly updated resource will help researchers and diagnosticians get a broad perspective of the most widely used - as well as the rare - biomarkers for COVID-19. The resource also encompasses risk biomarkers that will help the research and medical community stay abreast of the extensive research being conducted in the face of the ongoing pandemic.

## Conclusion

There is an urgent need within the research community to have an integrated and harmonized COVID-19 Biomarker resource. Variations in symptoms have not only exacerbated the diagnosis, prognosis and monitoring of COVID-19, but have also made it difficult to identify and develop vaccines and drugs. This resource has drawn from our extensive experience in integrating large biomarker (OncoMX - https://www.oncomx.org/) and glycoprotein (Glygen - https://www.glygen.org/) datasets to prepare a repository for COVID-19 biomarkers. As such, it currently includes over 500 biomarker entries, that encompassing both vastly studied and largely cross-referenced biomarkers, as well as rare and risk biomarkers. An overview of the current repository shows that the COVID-19 research community around the world is focused on the effects of the infection on the patient’s immune system, and various coagulopathies. Risk biomarkers included in our resource also shed light on at-risk patient populations and allows investigation of underlying comorbidities that might affect prognosis and treatment outcome. To further develop and improve the COVID-19 Biomarker resource, we envision (1) extending the search for new biomarkers in future curation rounds with adjusted query terms and parameters; (2) extending the data model and further data analyses to find possible correlations of biomarkers with patient features; and (3) examining possible multivariate biomarker profiles. To these ends, we recognize the need for standardization and formalization of the collected information in an ontology that connects biomarkers with their indications. Construction of such an ontology is underway. Nonetheless, in these early stages of understanding the pathology of this infectious disease, we provide this biomarker resource to support continued research around the world to better understand and manage COVID-19. Collective analyses of these biomarkers using a resource such as ours will help researchers gain a wider perspective of the disease state, with potential positive clinical impact. We encourage feedback and welcome contributions to the resource at https://data.oncomx.org/covid19.

## License

All data is freely available under the Creative Commons CC0 or CC-BY-4.0 license.

## Acknowledgements

Research reported in this article was supported in part by National Cancer Institute Grant No. U01CA215010 to RM.

HIVE Lab (https://hive.biochemistry.gwu.edu)

Jiuge Yang (content developer); Research Assistant

Sneh Talwar (content developer); Research Assistant

<u>Collaborators</u>

Dr. Shant Ayanyan (physician)

<u>Volunteers</u>

*Undergraduate students*

Miguel Mazumder (bioinformatics database Q/A support and curator)

Antarjot Kaur (bioinformatics curator)

Ruqaia Al-Kohlany (bioinformatics curator)

Mariana Escalante (bioinformatics curator)

Chakshu Gandhi (bioinformatics curator)

*High-school students*

Rita Mazumder (Asst. coordinator and bioinformatics curator)

Renee Long (bioinformatics curator)

Sara Burr (bioinformatics curator)

Kristina Ayers (bioinformatics curator)

Niharika Chanda (bioinformatics curator)

Rishab Desai (bioinformatics curator)

Andy Cao (bioinformatics curator)

Sejal Singh (bioinformatics curator)

Pranav Mishra (bioinformatics curator)

Nikita Wagle (bioinformatics curator)

Noyanika Vattathara (bioinformatics curator)

Sahana Ramesh (bioinformatics curator)

Avery Ye (bioinformatics curator)

Chakshu Gandhi (bioinformatics curator)

Jonathan Ye (bioinformatics curator)

Pranav Mishra (bioinformatics curator)

Dia Jhaveri (bioinformatics curator)

Siddharth Krishnan (bioinformatics curator)

Arya Adake (bioinformatics curator)

Anders Gyllenhoff (bioinformatics curator)

